## Supplementary Figures for "The Interpretable Multimodal Machine Learning (IMML) framework reveals pathological signatures of distal sensorimotor polyneuropathy"

**Supplementary Information**

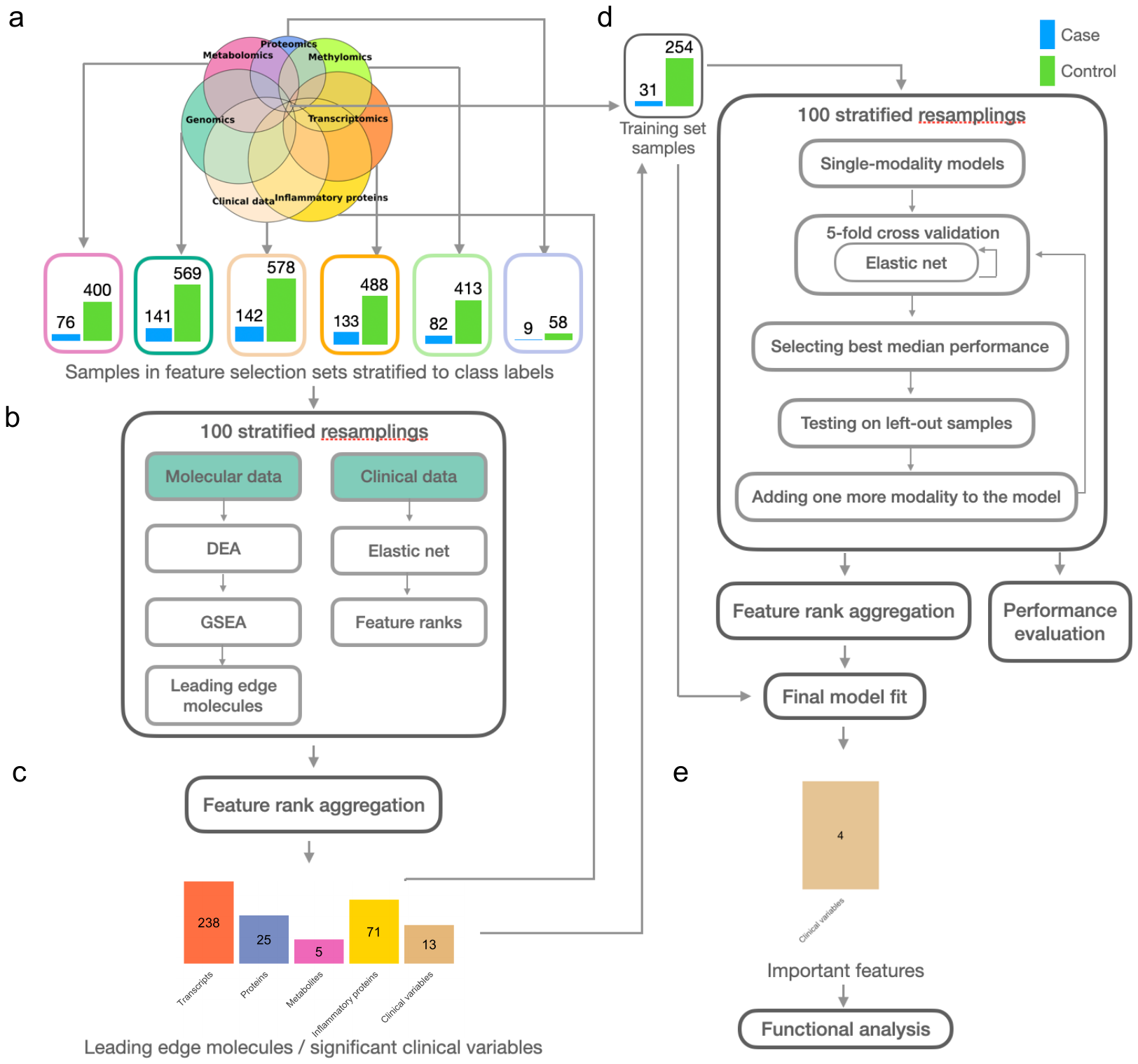

**Supplementary Figure S1. Analysis pipeline and results for cross-sectional DSPN**

a. Features in modality-specific datasets were selected independently using non-overlapping modality-specific samples. The selected features stratified into case and control are shown in the barplots.
b. Molecular data went through differential expression analysis (DEA) which generated a molecule list sorted by t-statistics which was then used as input for gene set enrichment analysis (GSEA). GSEA output leading edge genes which drive the enrichment of their respective gene sets. Clinical features was selected by training elastic net models and extracting important features. The process was repeated using 100 stratified resamplings.
c. The final significant list of molecules and clinical variables were selected using a rank aggregation algorithm.
d. After feature selection step, the selected features were then integrated to train models to predict DSPN, using the left-out overlapping dataset (training set). The training aimed to determine the optimal complexity and composition of the models by implementing elastic net with forward feature selection in a nested cross-validation manner, using weighted log loss as performance metric to account for class imbalance. We used 100 stratified resamplings during training and the rank aggregation at the end to select the most stable model.

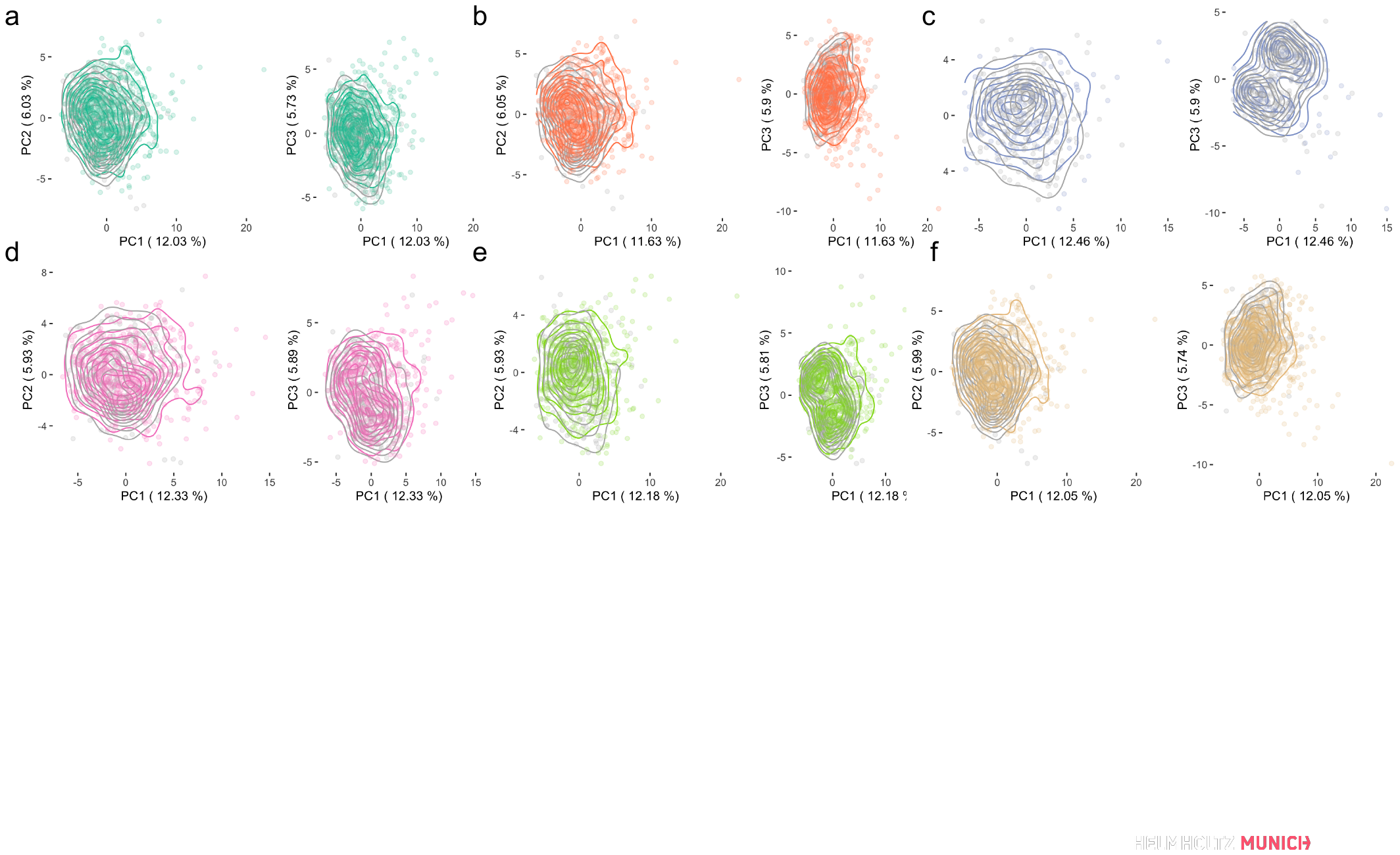

**Supplementary Figure S2.** PCA of clinical features for feature selection and model training datasets. Grey contour plots highlight the model training sets, whilst other colors indicate the feature selection set of the different modalities: (a) Genomics, (b) Transcriptomics, c) Proteomics, (d) Metabolomics, (e) Methylomics and (f) Clinical data.

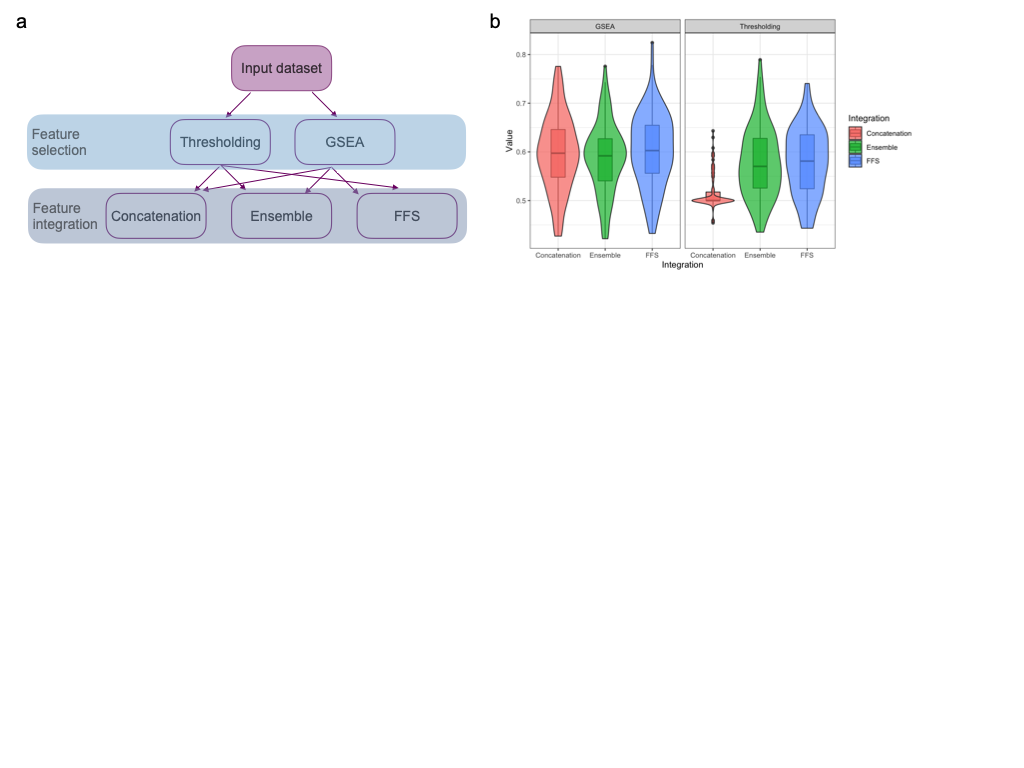

**Supplementary Figure S3. Benchmarking of feature selection and integration methods**

a. Illustration of methods for feature selection (thresholding and GSEA) and feature integration (concatenation, ensemble and our FFS algorithm) in a conventional multi-modal machine learning process. Arrows show the possible trajectory of the process in which different combinations of these methods could be used.
b. Benchmarking result showing prediction performance on the test set of different selection-integration methods for incident DSPN prediction using transcriptomic, proteomic, metabolomic and clinical data. Distributions of AUROC for the matched 100 stratified resamplings are shown in the y-axis and different methods are shown on the x-axis.

**Supplementary Figure S4. Network of enriched gene sets in cross-sectional DSPN**

Network of enriched gene sets from which the predictive features were selected, for cross-sectional DSPN prediction. Nodes are the gene sets coloured with their corresponding data modality. Size of the nodes reflects their centrality with respect to the network. Edges are the number of shared leading-edge molecules between two nodes.

**Supplementary Figure S5. Network of enriched features in cross-sectional DSPN**

Network of all selected features for training cross-sectional DSPN models. Nodes are the features coloured with their corresponding data modality. Edges are the number of shared gene sets between two nodes.

**Supplementary Figure S6. Network of enriched gene sets in incident DSPN**

Network of enriched gene sets from which the predictive features were selected, for incident DSPN prediction. Nodes are the gene sets coloured with their corresponding data modality. Size of the nodes reflects their centrality with respect to the network. Edges are the number of shared leading-edge molecules between two nodes.

**Supplementary Figure S7. Network of enriched features in incident DSPN**

Network of all selected features for training incident DSPN models. Nodes are the features coloured with their corresponding data modality. Edges are the number of shared gene sets between two nodes.

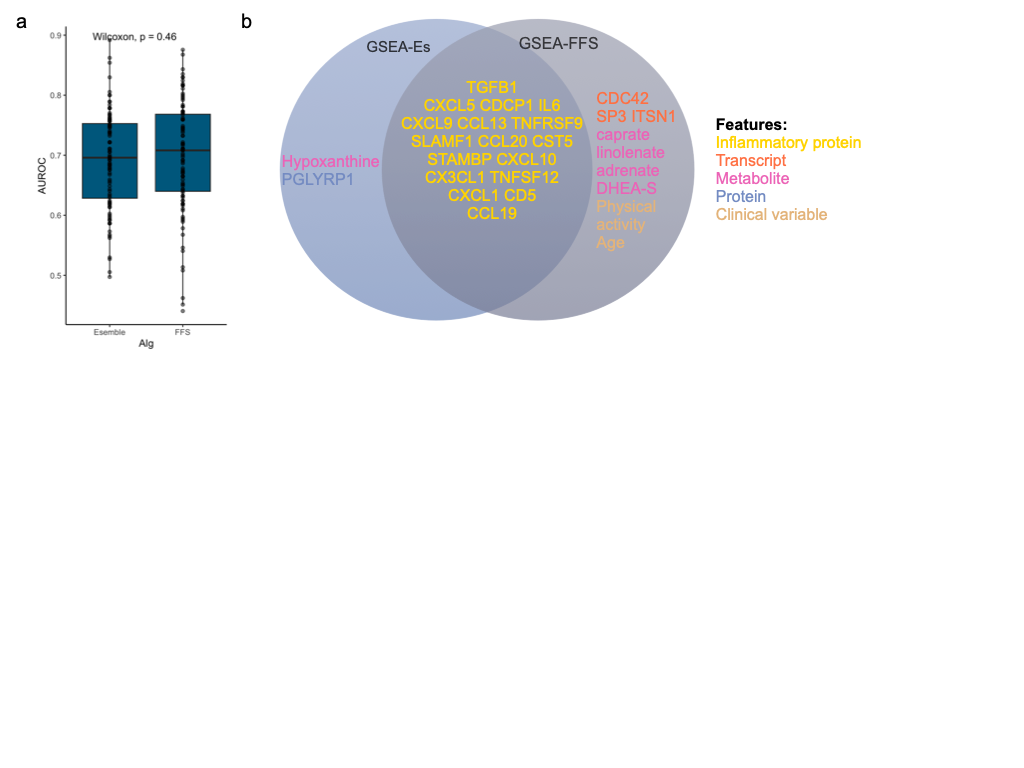

**Supplementary Figure S8. Performance of forward feature selection (FFS) and ensemble stacking feature integration methods across 100 stratified resamples.** (a) AUROC of the testing prediction of the two algorithms. P-value of Wilcoxon rank sum test is shown.(b) Important features selected by the GSEA-ensemble stacking (GSEA-Es) and GSEA-FFS methods and their overlapping.

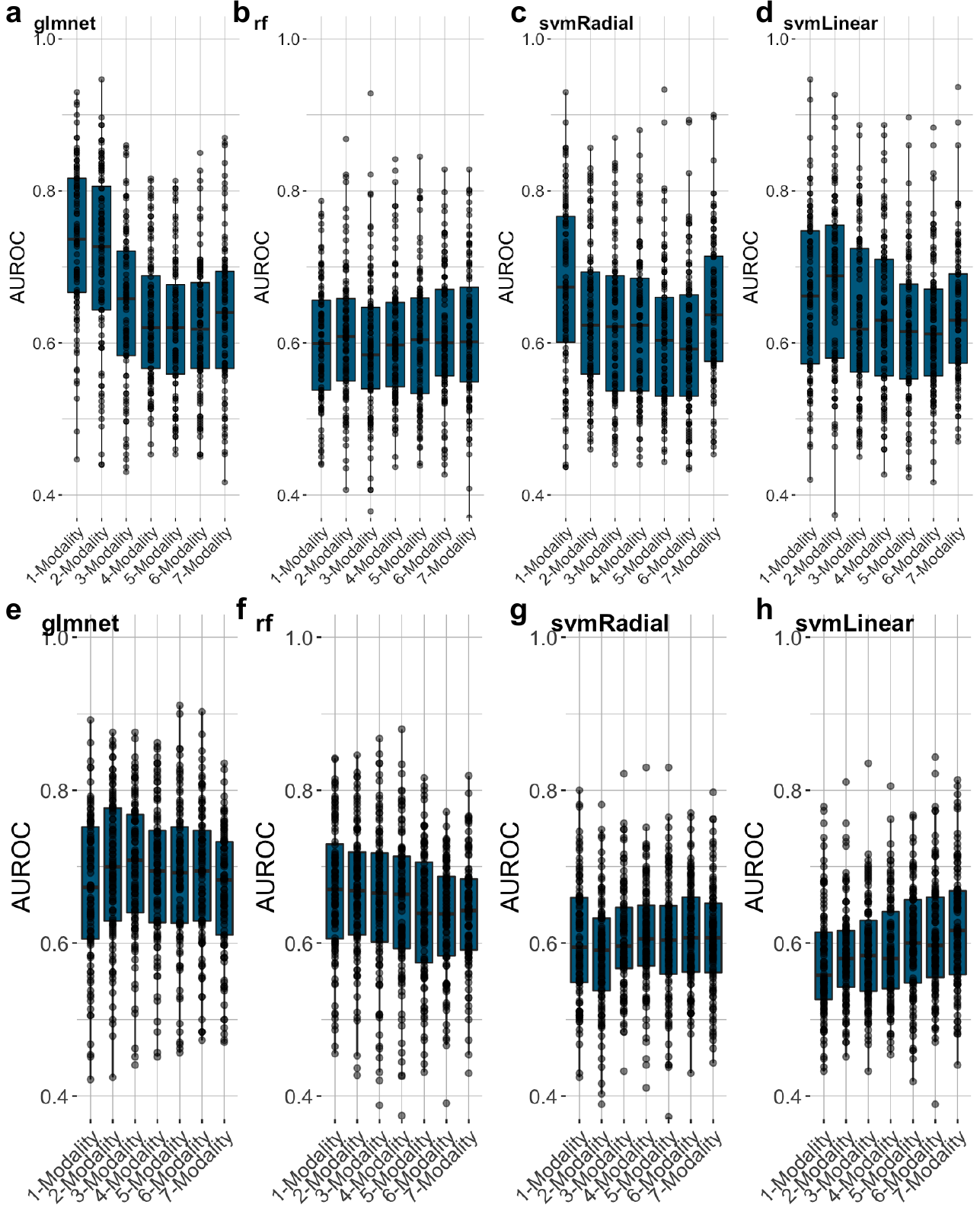

**Supplementary Figure S9: Prediction performance of four different machine learning algorithms.** Here we compare the predictive power of (a-d) prevalent DSPN and (e-h) incident DSPN. We benchmarked (a,e) elastic net (glmnet), (b,f) random forest (rf), and support vector machine with (c,g) radial (svmRadial) and (d,h) linear kernel (svmLinear).

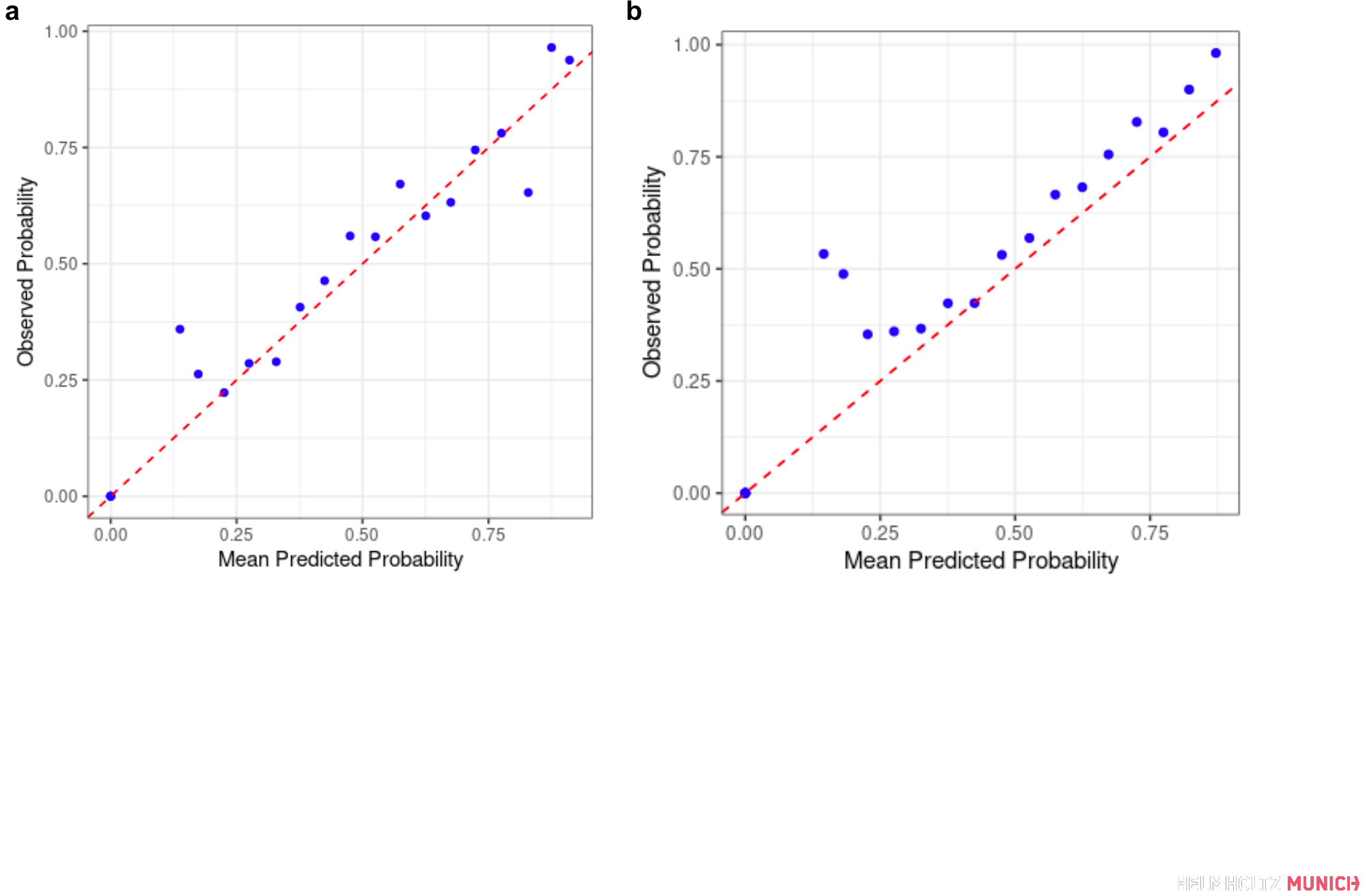

**Supplementary Figure S10: Calibration plots of predicted probabilities for prevalent DSPN (a) and incident DSPN (b).** The predicted probabilities were calibrated using the Platt scaling method.

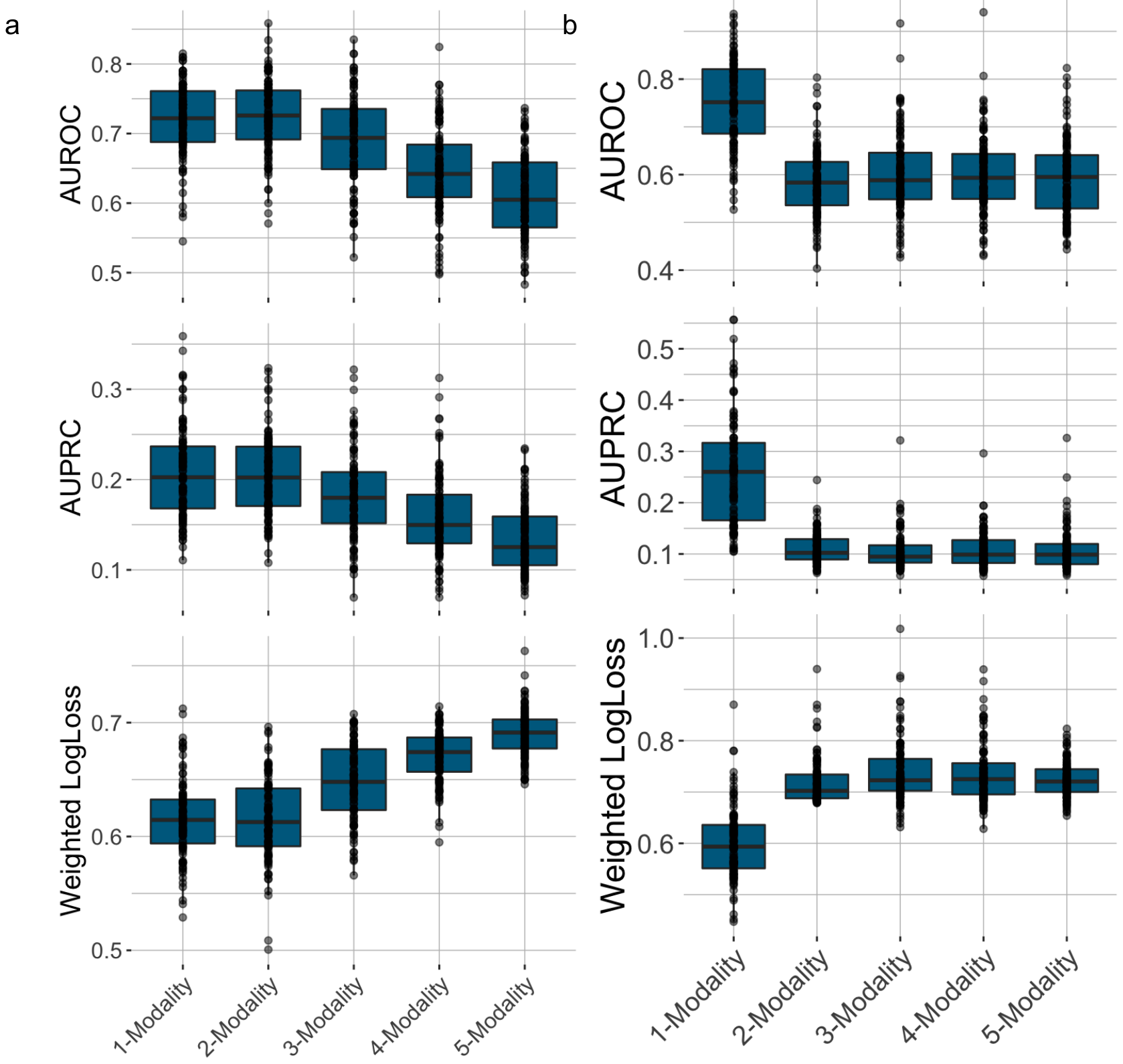

**Supplementary Figure S11. Prediction performance of prevalent DSPN models, when forcing the FFS algorithm to choose clinical model at the beginning.**

a. Prediction performance during cross-validation. X-axis shows the increasing model complexity. Y-axis shows the median of performance values across 5-fold cross-validation for AUROC, AUPRC and weighted log-loss
b. Prediction performance on the testing sets. X-axis shows the increasing model complexity. Y-axis shows the performance values on the testing sets for AUROC, AUPRC and weighted log-loss

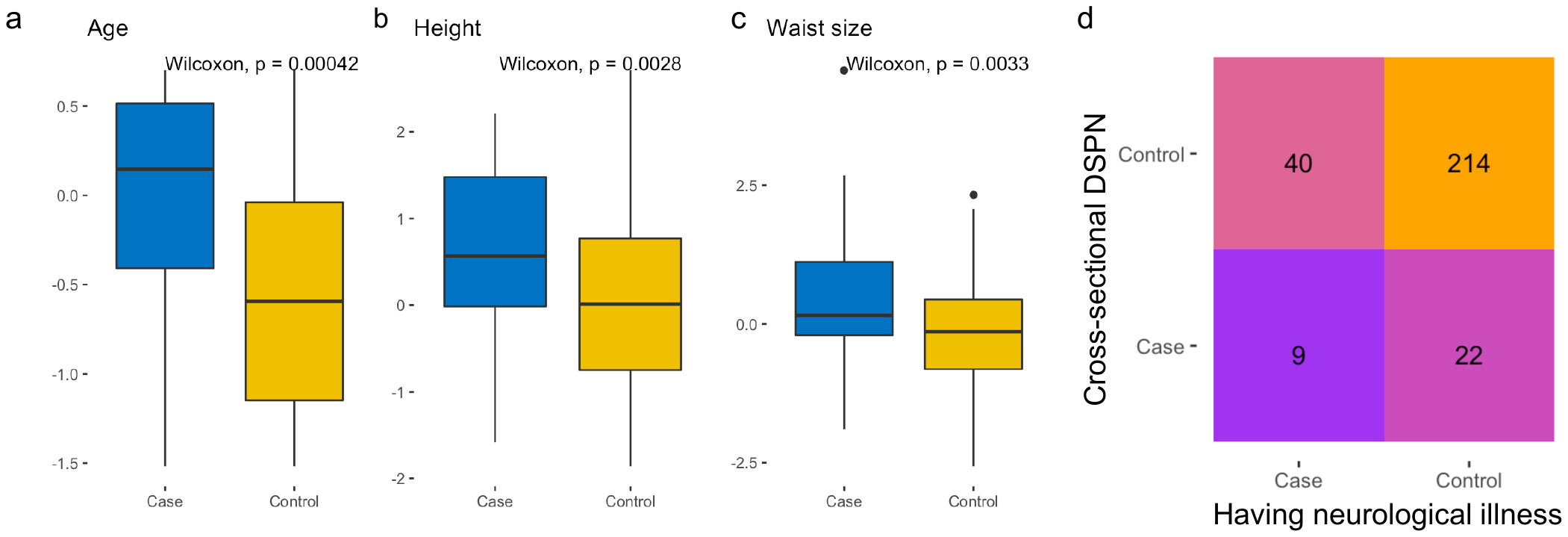

**Supplementary Figure S12. Distribution of important clinical variables for cross-sectional DSPN model**

Distribution of age, height and waist size in the training set stratified into case and control (panel a, b and c respectively). Panel d shows association of patients who have neurological illness in general and cross-sectional DSPN. P-values for Wilcoxon rank sum test and Fisher’s exact test are shown.

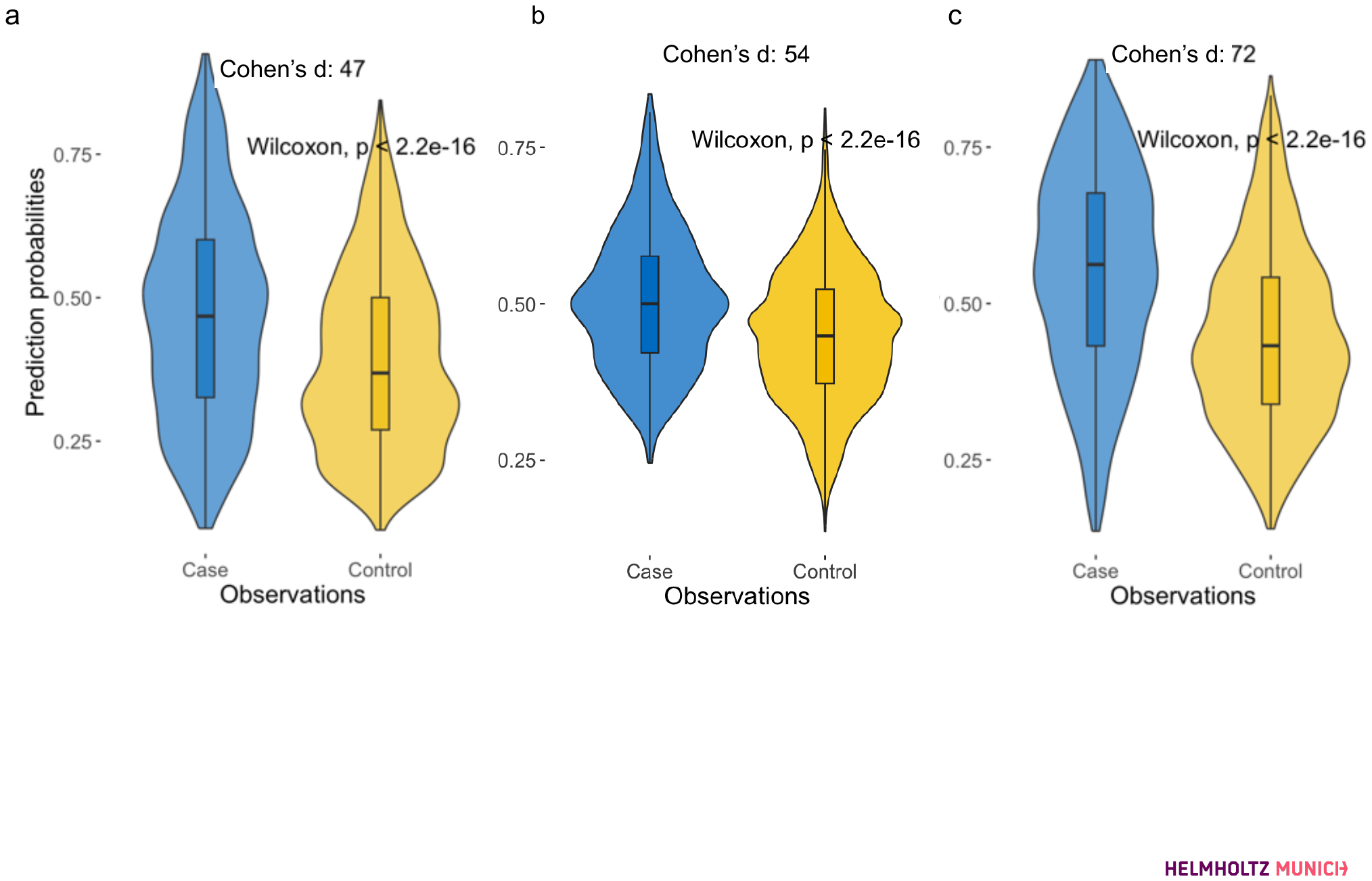

**Supplemental Figure S13: Baseline models to predict DSPN incidence**. Prediction probabilities during testing of negative samples using a) the prevalent DSPN model trained on clinical data alone at F4, b) baseline incidence model trained only on clinical variables at F4 and incidence label at FF4 and c) the full incidence model trained on clinical + molecular variables at F4 and incidence label at FF4. Cases are samples developing DSPN from F4 to FF4, and controls are ones remaining negative. For each comparison, Cohen’s d was used as the measure of the difference between groups.

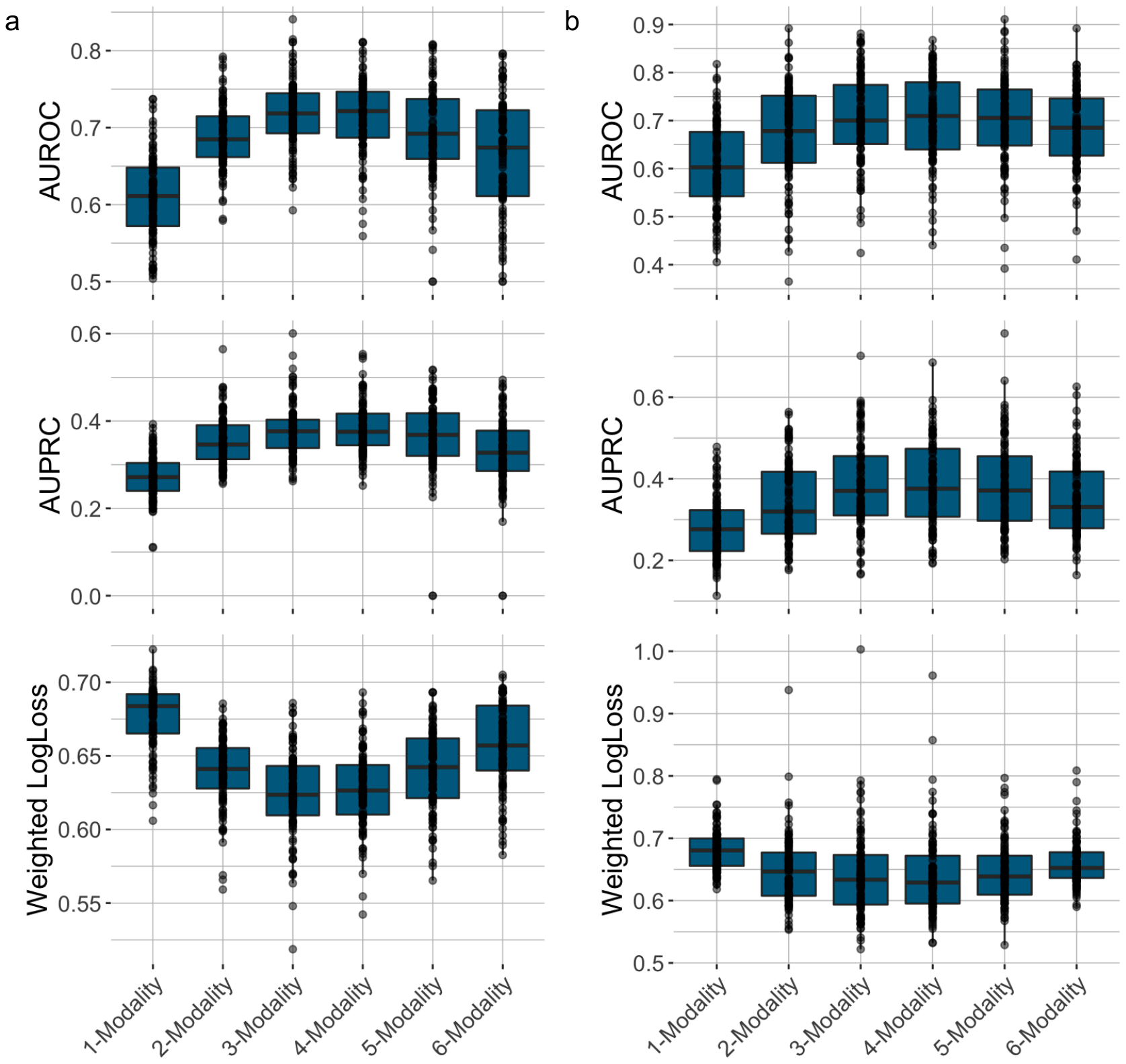

**Supplementary Figure S14. Prediction performance of incident DSPN models, when forcing the FFS algorithm to choose clinical model at the beginning.**

a. Prediction performance during cross-validation. X-axis shows the increasing model complexity. Y-axis shows the median of performance values across 5-fold cross-validation for AUROC, AUPRC and weighted log-loss
b. Prediction performance on the testing sets. X-axis shows the increasing model complexity. Y-axis shows the performance values on the testing sets for AUROC, AUPRC and weighted log-loss

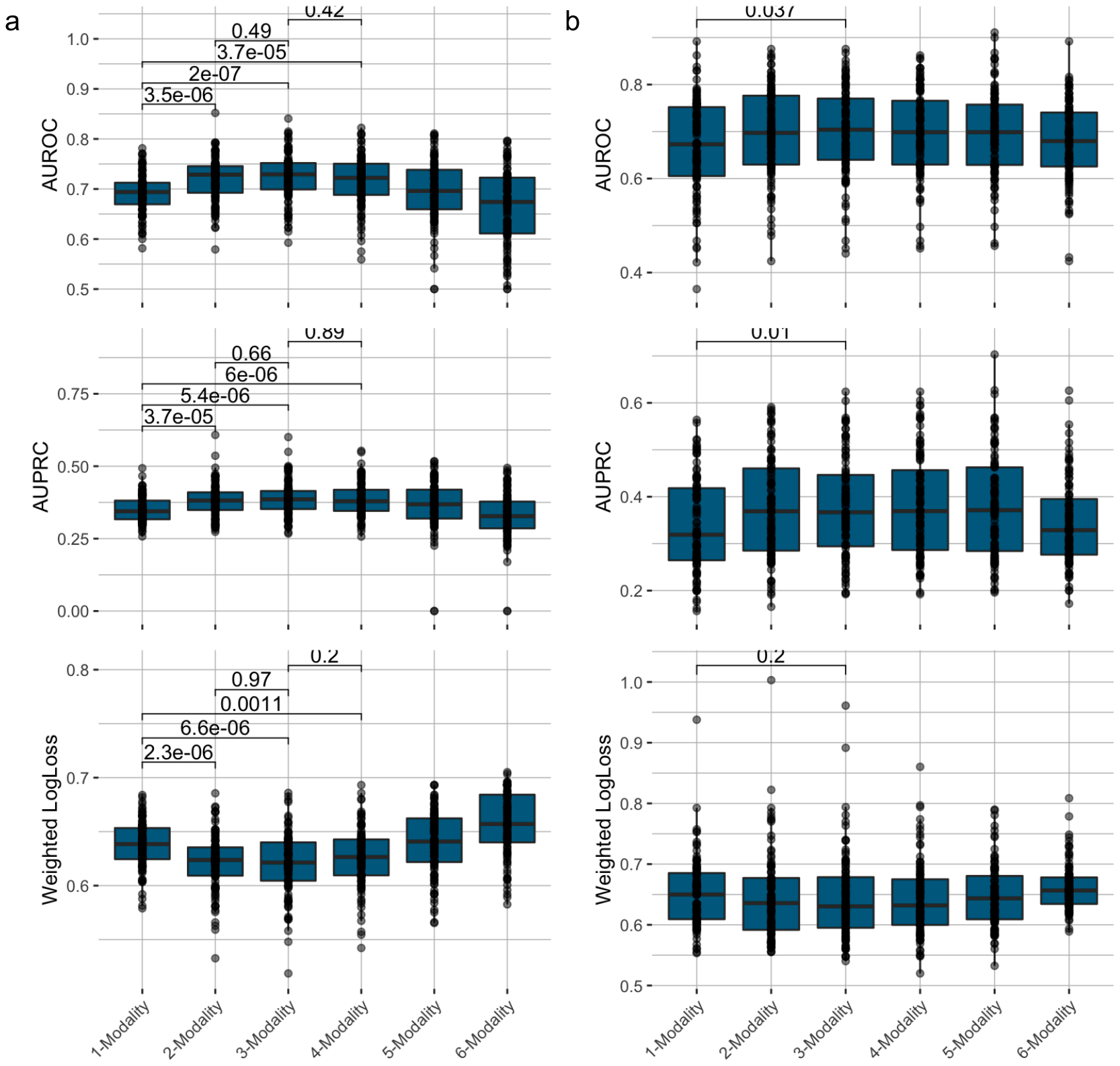

**Supplementary Figure S15. Prediction performance of incident DSPN models, when allowing the FFS algorithm to choose starting model based on cross-validation.**

a. Prediction performance during cross-validation. X-axis shows the increasing model complexity. Y-axis shows the median of performance values across 5-fold cross-validation for AUROC, AUPRC and weighted log-loss
b. Prediction performance on the testing sets. X-axis shows the increasing model complexity. Y-axis shows the performance values on the testing sets for AUROC, AUPRC and weighted log-loss

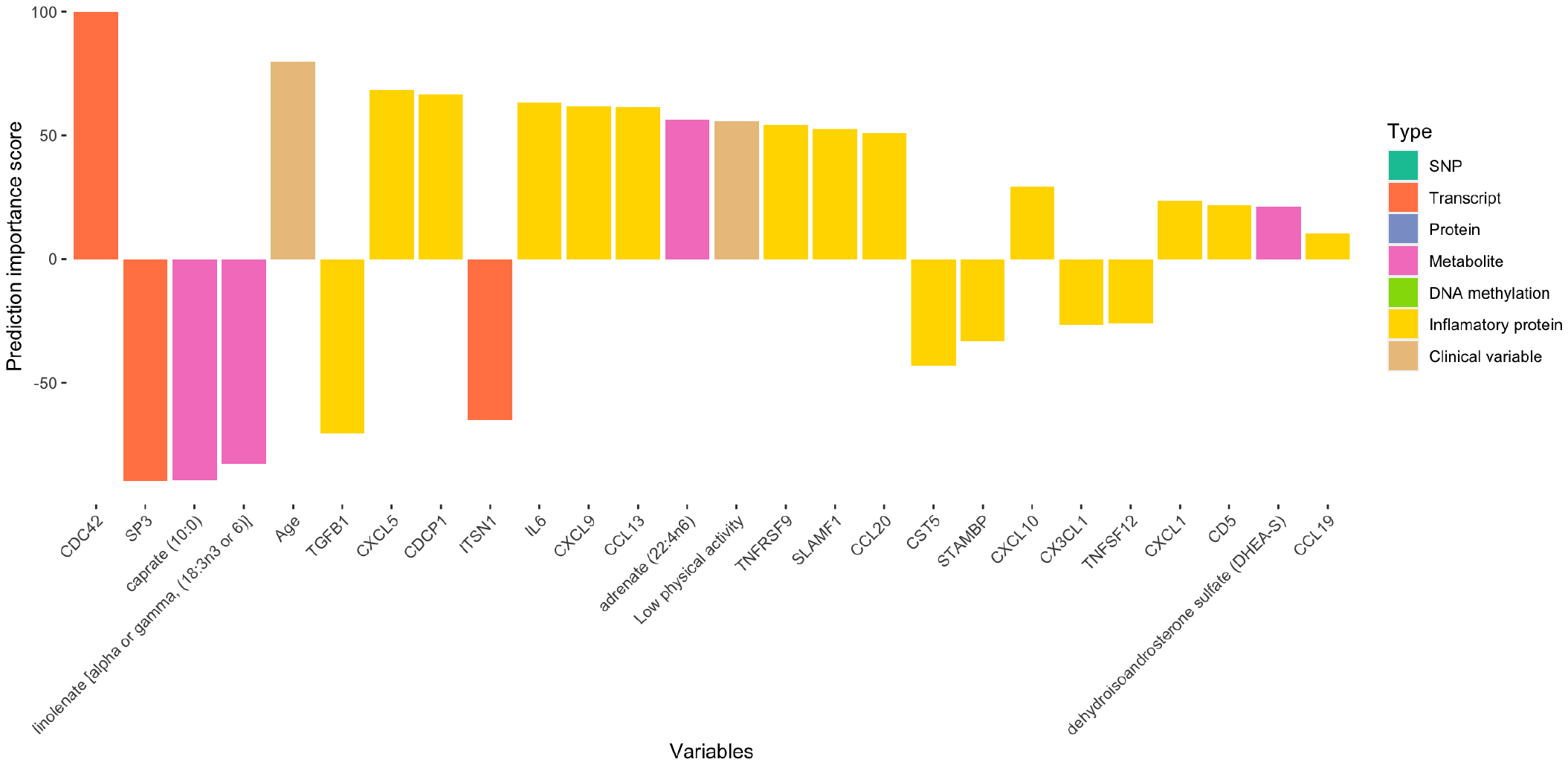

**Supplementary Figure S16. Feature importance score of the important features of the final incident DSPN model**

X-axis shows the features in decreasing magnitude of the t-statistics in the final model. Y-axis shows the t-statistics (signed importance scores) of the features. Colors represent the data modality.

**Supplementary Figure S17. Distribution of important clinical variables for incident DSPN model**

Distribution of the features in the training set stratified into case and control. P-values for Wilcoxon rank sum test and Fisher’s exact test are shown.

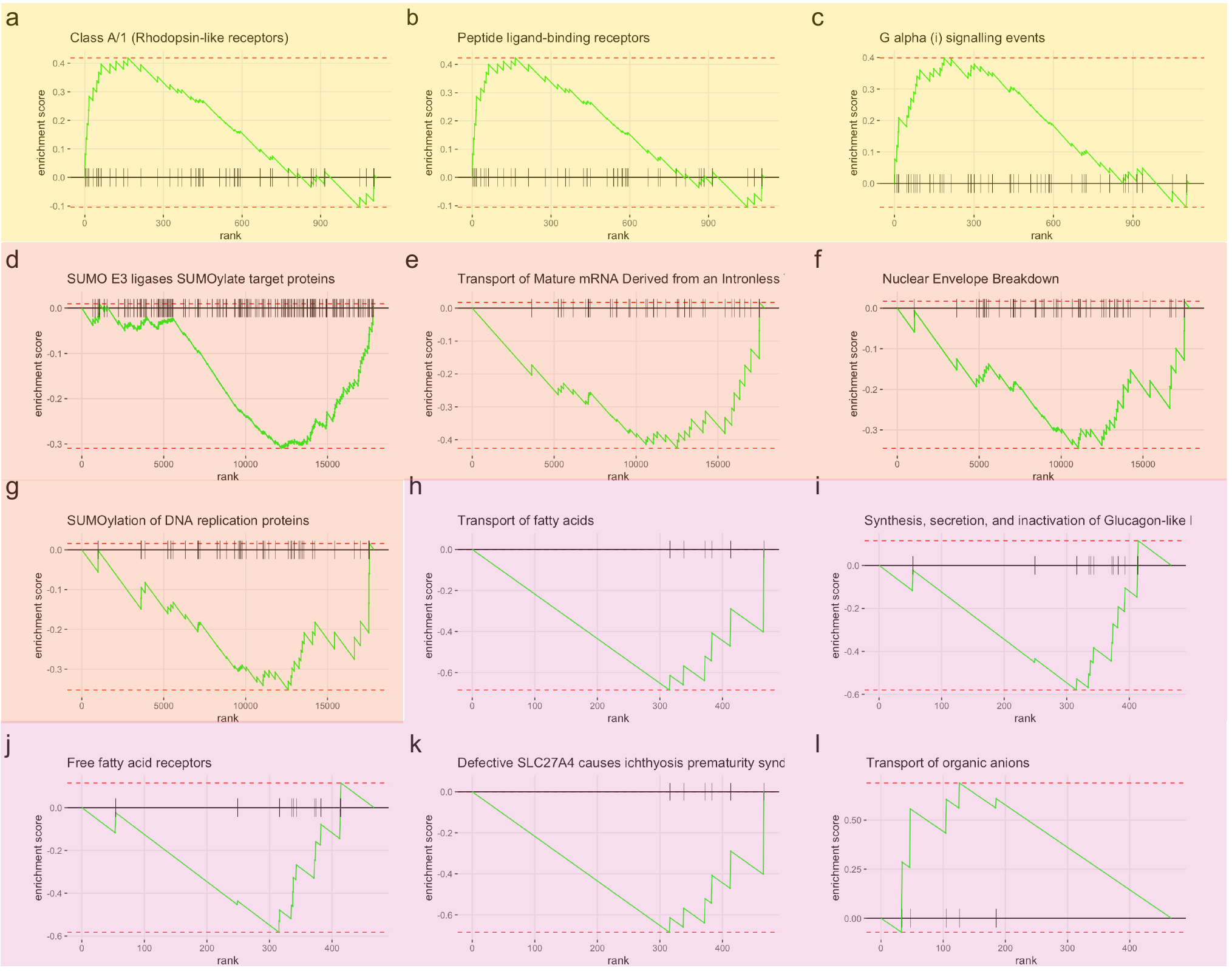

**Supplementary Figure S18. Examples of consistently enriched signalling pathways that are predictive of incident DSPN**

X-axis represents all evaluated genes ranked in decreasing order of t-statistics, with ticks represent genes that belong to the examined gene set. Y-axis represent the enrichment score. Panels a-c are inflammation protein pathways, d-g are transcriptomic pathways and h-l are metabolomic pathways.

**Supplementary Table S1. Clinical characteristics of the dataset for prevalent DSPN prediction**

| **Variable** | **Control (MNSI <3)** | **Case (MNSI >= 3)** | **P** |
| --- | --- | --- | --- |
| N | 903 | 188 |  |
| Age, years | 69.7 ± 5.2 | 72.5 ± 5.2 | 1.09e-10 |
| Sex, % male | 49.4 | 60.6 | 0.005 |
| Height, cm | 165.3 ± 8.8 | 167.9 ± 9.6 | 0.00071 |
| BMI, kg/m2 | 28.4 ± 4.2 | 30.2 ± 5.2 | 1.30e-05 |
| Waist circumference, cm | 97.2 ± 11.7 | 103.7 ± 12.9 | 8.11e-10 |
| Systolic blood pressure, mmHg | 128.8 ± 20 | 128.6 ± 20 | 0.873 |
| Diastolic blood pressure, mmHg | 74.4 ± 10.1 | 72.4 ± 9.8 | 0.007 |
| Hypertension, % | 62.0 | 64.4 | 0.561 |
| Smoking, %, never/former/current | 51.6/40.7/7.7 | 44.9/48.1/7.0 | 0.233 |
| High alcohol consumption, % | 29.1 | 33.7 | 0.220 |
| Low physical activity, % | 36.8 | 51.9 | 0.014 |
| Previous myocardial infarction, % | 5.9 | 9.1 | 0.104 |
| Previous stroke, % | 3.2 | 8.0 | 0.006 |
| Presence of neurological diseases, % | 16.2 | 31.0 | 4.33e-06 |
| Absent ankle reflexes, % | 5 | 72.3 | 6.63e-112 |
| Foot ulcer present, % | 0 | 2.1 | 0.001 |
| MNSI score | 1.7 ± 1 | 4.3 ± 0.9 | 2.34e-107 |
| Use of NSAIDs, % | 3.4 | 7.4 | 0.024 |
| NGT, % | 53.7 | 45.7 | 0.054 |
| i-IFG, % | 5.3 | 3.7 | 0.464 |
| i-IGT, % | 16.7 | 12.2 | 0.154 |
| IFG/IGT, % | 4.3 | 6.9 | 0.133 |
| Newly diagnosed diabetes, % | 6.4 | 4.8 | 0.504 |
| Known diabetes, % | 13.5 | 26.6 | 1.25e-05 |
| Diabetes duration, years* | 8.1 ± 6.4 | 15 ± 10.6 | 1.58e-15 |
| Metabolic parameters |  |  |  |
| Fasting glucose, mg/dL^+^ | 103.6 ± 21.2 | 110.4 ± 29.9 | 0.015 |
| 2-h glucose, mg/dL^+^ | 128.0 ± 41.9 | 127.2 ± 38.6 | 0.945 |
| HbA1c, % | 5.7 ± 0.7 | 6.0 ± 0.8 | 3.06e-06 |
| Total cholesterol, mg/dL | 222.7 ± 41.0 | 210.8 ± 37.9 | 0.00014 |
| LDL cholesterol, mg/dL | 140.7 ± 36.2 | 131.7 ± 33.4 | 0.001 |
| HDL cholesterol, mg/dL | 56.0 ± 14.3 | 53.4 ± 12.2 | 0.075 |
| Creatinine, mg/dL | 0.95 ± 0.3 | 1.02 ± 0.3 | 0.001 |
| Uric acid, mg/dL | 5.5 ± 1.4 | 5.8 ± 1.5 | 0.015 |
| * Only applicable to people with diabetes |  |  |  |
| + Only applicable to people without known diabetes | |  |  |

**Supplementary Table S2. Clinical characteristics of the dataset for incident DSPN prediction**

| **Variable** | **Control (no incident F4->FF4)** | **Case (incident F4-> FF4)** | **P** |
| --- | --- | --- | --- |
| N | 394 | 131 |  |
| Age, years | 68.0 ± 4.6 | 70.1 ± 4.9 | 2.46e-05 |
| Sex, % male | 49.2 | 56.5 | 0.159 |
| Height, cm | 165.9 ± 8.5 | 167.6 ± 9.4 | 0.064 |
| BMI, kg/m2 | 27.7 ± 3.8 | 29.1 ± 4.0 | 0.00054 |
| Waist circumference, cm | 94.8 ± 11.2 | 99.9 ± 11.4 | 1.34e-05 |
| Systolic blood pressure, mmHg | 128.4 ± 19.2 | 131.3 ± 19.9 | 0.217 |
| Diastolic blood pressure, mmHg | 75.5 ± 10.1 | 75.5 ± 9.2 | 0.950 |
| Hypertension, % | 56.3 | 65.6 | 0.066 |
| Smoking, %, never/former/current | 52.0/42.4/5.6 | 55.0/33.6/11.4 | 0.054 |
| High alcohol consumption, % | 29.4 | 35.9 | 0.191 |
| Low physical activity, % | 26.4 | 42.7 | 0.00064 |
| Previous myocardial infarction, % | 4.8 | 6.9 | 0.373 |
| Previous stroke, % | 1.0 | 0.8 | 1 |
| Presence of neurological diseases, % | 14.7 | 21.4 | 0.102 |
| Absent ankle reflexes, % | 3.8 | 6.1 | 0.323 |
| Foot ulcer present, % | 0 | 0 | 1 |
| MNSI score | 1.5 ± 1.0 | 1.9 ± 0.9 | 2.65e-05 |
| Use of NSAIDs, % | 1.0 | 2.3 | 0.374 |
| NGT, % | 62.9 | 50.4 | 0.013 |
| i-IFG, % | 3.0 | 7.6 | 0.040 |
| i-IGT, % | 14.5 | 16.8 | 0.573 |
| IFG/IGT, % | 4.6 | 4.6 | 1 |
| Newly diagnosed diabetes, % | 5.6 | 5.3 | 1 |
| Known diabetes, % | 9.4 | 15.3 | 0.074 |
| Diabetes duration, years* | 6.9 ± 5.5 | 8.9 ± 5.2 | 0.116 |
| Metabolic parameters |  |  |  |
| Fasting glucose, mg/dL^+^ | 101.0 ± 16.4 | 103.8 ± 17.2 | 0.078 |
| 2-h glucose, mg/dL^+^ | 123.9 ± 38.6 | 127.4 ± 38.4 | 0.371 |
| HbA1c, % | 5.7 ± 0.5 | 5.8 ± 0.7 | 0.027 |
| Total cholesterol, mg/dL | 226.3 ± 40.5 | 216.1 ± 42.7 | 0.009 |
| LDL cholesterol, mg/dL | 142.5 ± 36.3 | 136.6 ± 37.5 | 0.069 |
| HDL cholesterol, mg/dL | 57.3 ± 14.2 | 52.5 ± 12.3 | 0.00025 |
| Creatinine, mg/dL | 0.9 ± 0.2 | 1.0 ± 0.3 | 0.071 |
| Uric acid, mg/dL | 5.5 ± 1.3 | 5.6 ± 1.4 | 0.692 |
| * Only applicable to people with diabetes | |  |  |
| + Only applicable to people without known diabetes | |  |  |

**Supplementary Table S3. Significantly enriched signalling pathways during feature selection for prevalent DSPN prediction**

| **pathway** | **pval** | **padj** | **ES** | **NES** | **size** | **Type** |
| --- | --- | --- | --- | --- | --- | --- |
| Formation of a pool of free 40S subunits | 1.874e-06 | 2.024e-05 | -0.445 | -2.140 | 95 | Transcriptomics |
| GTP hydrolysis and joining of the 60S ribosomal subunit | 3.308e-05 | 8.132e-05 | -0.405 | -1.986 | 104 | Transcriptomics |
| L13a-mediated translational silencing of Ceruloplasmin expression | 3.555e-05 | 8.132e-05 | -0.404 | -1.972 | 103 | Transcriptomics |
| Nonsense Mediated Decay (NMD) enhanced by the Exon Junction Complex (EJC) | 6.2356e-05 | 0.0001 | -0.395 | -1.944 | 106 | Transcriptomics |
| Collagen chain trimerization | 0.0001 | 0.0002 | 0.522 | 2.029 | 39 | Transcriptomics |
| Influenza Infection | 0.0002 | 0.0004 | -0.341 | -1.762 | 143 | Transcriptomics |
| Collagen biosynthesis and modifying enzymes | 0.0003 | 0.0005 | 0.441 | 1.921 | 61 | Transcriptomics |
| Assembly of collagen fibrils and other multimeric structures | 0.0003 | 0.0005 | 0.448 | 1.928 | 58 | Transcriptomics |
| Degradation of the extracellular matrix | 0.0004 | 0.0006 | 0.338 | 1.678 | 131 | Transcriptomics |
| Formation of the ternary complex and subsequently the 43S complex | 0.0005 | 0.0007 | -0.460 | -1.941 | 47 | Transcriptomics |
| SUMOylation of DNA methylation proteins | 0.0015 | 0.002 | -0.635 | -2.006 | 15 | Transcriptomics |
| Selenoamino acid metabolism | 0.0018 | 0.002 | -0.333 | -1.642 | 110 | Transcriptomics |
| Major pathway of rRNA processing in the nucleolus and cytosol | 0.0019 | 0.002 | -0.293 | -1.550 | 170 | Transcriptomics |
| Ribosomal scanning and start codon recognition | 0.002 | 0.003 | -0.413 | -1.774 | 53 | Transcriptomics |
| Regulation of expression of SLITs and ROBOs | 0.005 | 0.005 | -0.291 | -1.526 | 159 | Transcriptomics |
| Laminin interactions | 0.0001 | 0.0713 | 0.829 | 2.017 | 9 | Proteomics |
| Antimicrobial peptides | 0.0004 | 0.0713 | -0.676 | -2.111 | 16 | Proteomics |
| Interleukin-20 family signaling | 0.0004 | 0.0713 | 0.745 | 2.009 | 13 | Proteomics |
| Interleukin-3 Interleukin-5 and GM-CSF signaling | 0.0006 | 0.082 | 0.618 | 1.922 | 23 | Proteomics |
| Transport of nucleosides and free purine and pyrimidine bases across the plasma membrane | 0.0002 | 0.0002 | -0.931 | -2.083 | 5 | Metabolomics |
